# Evaluation of Reproducibility in Urology Publications

**DOI:** 10.1101/773945

**Authors:** Shelby Lynn Rauh, Bradley S. Johnson, Aaron Bowers, Daniel Tritz, Benjamin Matthew Vassar

**Author notes:** **Corresponding Author:** Mrs. Shelby Lynn Rauh, Oklahoma State University Center for Health Sciences, 1111 W 17th St., Tulsa, OK 74107, USA.

## Abstract

**Take Home Message:** Many components of transparency and reproducibility are lacking in urology publications, making study replication, at best, difficult.

**Introduction:** Reproducibility is essential for the integrity of scientific research. Reproducibility is measured by the ability of investigators to replicate the outcomes of an original publication by using the same materials and procedures.

**Methods:** We sampled 300 publications in the field of urology for assessment of multiple indicators of reproducibility, including material availability, raw data availability, analysis script availability, pre-registration information, links to protocols, and whether the publication was freely available to the public. Publications were also assessed for statements about conflicts of interest and funding sources.

**Results:** Of the 300 sample publications, 171 contained empirical data and could be analyzed for reproducibility. Of the analyzed articles, 0.58% (1/171) provided links to protocols, and none of the studies provided analysis scripts. Additionally, 95.91% (164/171) did not provide accessible raw data, 97.53% (158/162) did not provide accessible materials, and 95.32% (163/171) did not state they were pre-registered.

**Conclusion:** Current urology research does not consistently provide the components needed to reproduce original studies. Collaborative efforts from investigators and journal editors are needed to improve research quality, while minimizing waste and patient risk.

## Introduction

Reproducibility is determined by the availability of materials, raw data, analysis procedures, and protocols used to conduct original research so that other researchers may replicate the findings; it is crucial to establishing credible and reliable research that ultimately governs clinical practice. Recent evidence suggests that up to 90% of preclinical research may not be reproducible.[1] A recent survey of over 1500 researchers concurred with this assessment, with the vast majority believing that biomedical research is experiencing a “reproducibility crisis”.[2] Several explanations have been offered for why reproducibility has become an issue, with pressure to publish and the race to be the first to report new findings being among the most likely causes.[3] When research is not reproducible, time and money are wasted reproducing erroneous results, and patients may be exposed to ineffective or harmful therapies.[4] Concerns about reproducibility span from preclinical to clinical research.

The field of prostate cancer research serves as an example. On the diagnostic side, *in vitro* studies are performed on prostate biopsy samples to advance understanding of early detection and diagnosis. However, widespread misuse of immunohistochemical staining exists, which contributes to the lack of research reproducibility. Sfanos *et al[5]* argued that the ubiquitously used research-grade antibodies within the biomedical research community (as opposed to clinical grade used for patient diagnosis) are not routinely validated in the investigators’ laboratories, which may lead to variable results that cannot be reproduced in subsequent studies. On the other end of the research spectrum, randomized clinical trials are conducted to evaluate the efficacy of new therapeutic agents for prevention or treatment of prostate cancer. In one large-scale randomized trial, Thompson *et al[6]* compared the effects of finasteride against placebo for prostate cancer prevention. These investigators found that finasteride prevented or delayed the development of prostate cancer but also led to an increased risk of higher-grade cancer upon detection. The raw data from this clinical trial were not made entirely available because of patient privacy and data “messiness”. Some investigators attempted to reanalyze the trial data, but the results were mixed[7,8]. Since then, Baker *et al[9]* proposed a method to overcome issues of privacy and messiness, while also fostering reproducibility of the trial outcomes.

**Figure 1:**
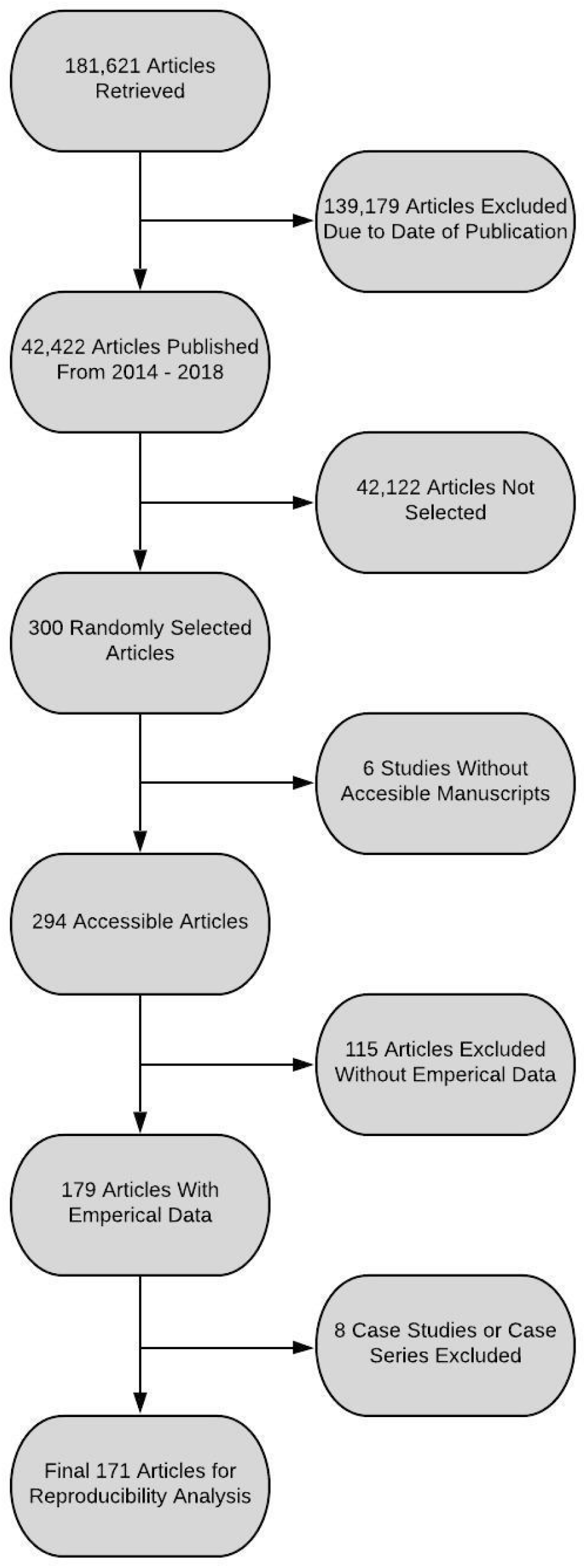
Flow Diagram of Included and Excluded Studies for the Reproducibility Analysis

Thus, when a study does not report the components needed to reproduce it, or when studies are not replicated by other researchers, determining the credibility of the original findings is hindered. Our study examines existing research in urology and determines how often studies include markers of reproducibility and how frequently studies are replicated. This research highlights the issue of reproducibility in urology, a field in which the topic has not been well explored. We anticipate that our findings will prompt discussions among investigators and journal editors, which may lead to improvement in the quality of research in the field.

## Methods

We used an observational, cross-sectional study design, drawing on the methodology of Hardwicke *et al[10]*, with modifications. This study did not involve human participants and was not subject to oversight or approval by an institutional review board.[11] We report our study in accordance with previously published guidelines for meta-epidemiological methodology research.[12] To foster transparency and reproducibility, we have uploaded our protocol, data extraction form, and other materials for public viewing on the Open Science Framework (OSF; https://osf.io/n4yh5/).

### Journal Selection

We used the National Library of Medicine (NLM) catalog to search for all relevant journals, using the subject terms tag Urology[ST]. The search was performed on 30 May 2019. The inclusion criteria required journals to have full-text publications in English and be MEDLINE indexed. The list of journals in the NLM catalog fitting the inclusion criteria were then extracted using the electronic International Standard Serial Number (ISSN) or the linking ISSN when the electronic ISSN was unavailable. PubMed was searched with the list of ISSNs to identify all articles published from 1 January 2014 to 31 December 2018. We randomly sampled 300 publications that met the inclusion criteria (https://osf.io/csf5t/).

### Data Extraction Training

The two investigators responsible for data extraction (S.R. and B.J.) underwent a full day of training to ensure adequate interrater reliability. The training included an in-person session to review the project study design, protocol, data extraction form, and examples of where information may be contained using two example publications. The investigators were then given three example publications from which to extract data in a blinded fashion. Afterward, the pair reconciled differences in their results. This training session was recorded from the presenter’s point of view (D.T.) and listed online for reference (https://osf.io/tf7nw/). As a final training exercise, investigators extracted data from the first 10 publications of the full sample and then met to reconcile any differences in the data before proceeding to extraction of the remaining 290 publications.

### Data Extraction

Data extraction on the remaining 290 publications was conducted in a duplicate, blinded fashion. A final consensus meeting was held with both investigators to resolve disagreements. A third investigator (D.T.) was available for adjudication but was not needed. Data were extracted using a pilot-tested Google form based on Hardwicke *et al*, with modifications.[10] This form contained information necessary for a study to be reproducible, such as the availability of materials, data, protocols, or analysis scripts (https://osf.io/3nfa5/). The data extracted varied based on the study design, with studies having no empirical data being excluded (e.g., editorials, commentaries [without reanalysis], simulations, news, reviews, and poems) (Table 1). The form also included the 5-year impact factor and that of the most recent year available and expanded the study design options to include cohort studies, case series, secondary analyses, chart reviews, and cross-sectional studies. Funding options were also expanded to include university, hospital, public, private/industry, non-profit, or mixed funding.

**Table 1:**
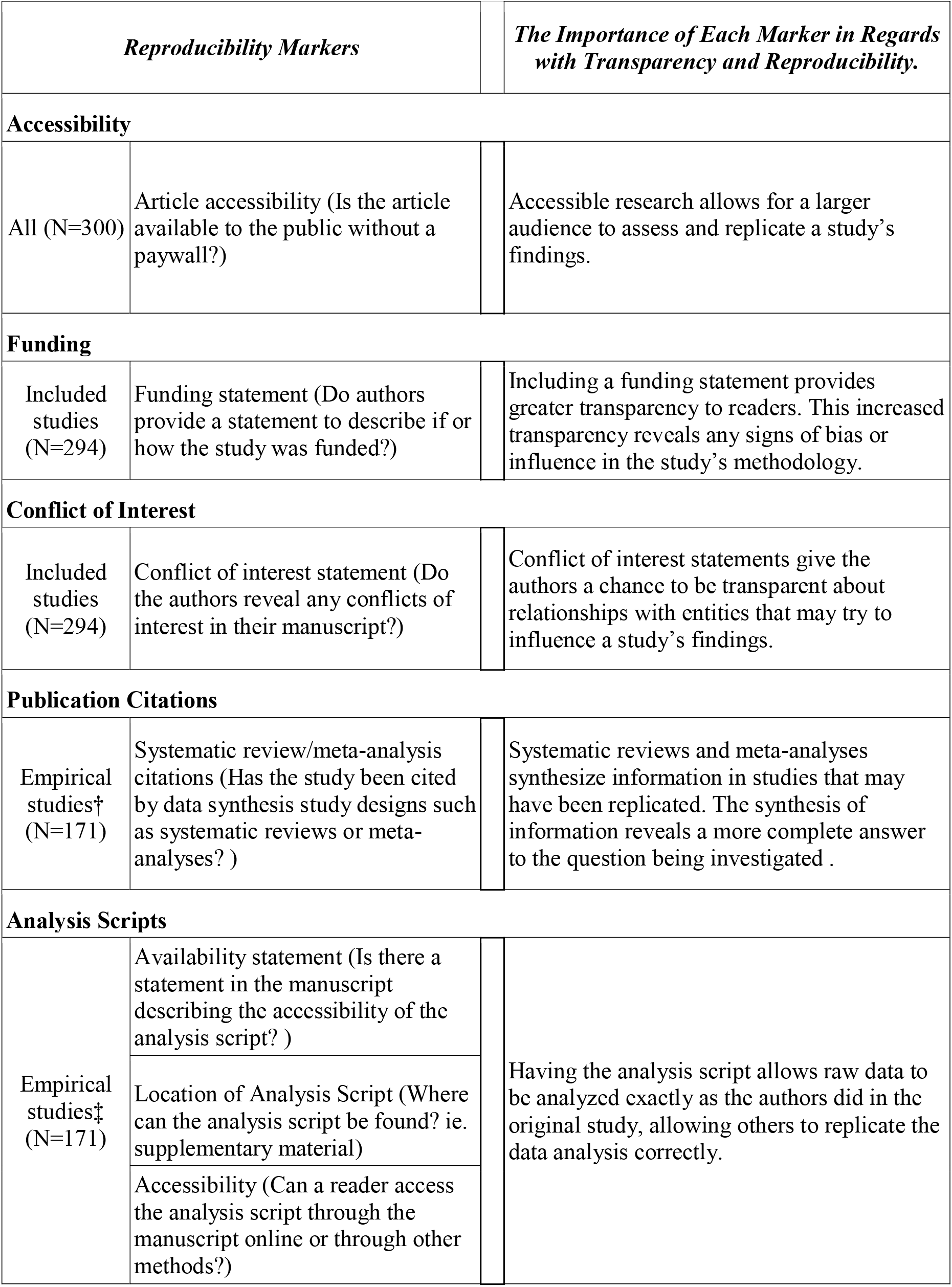
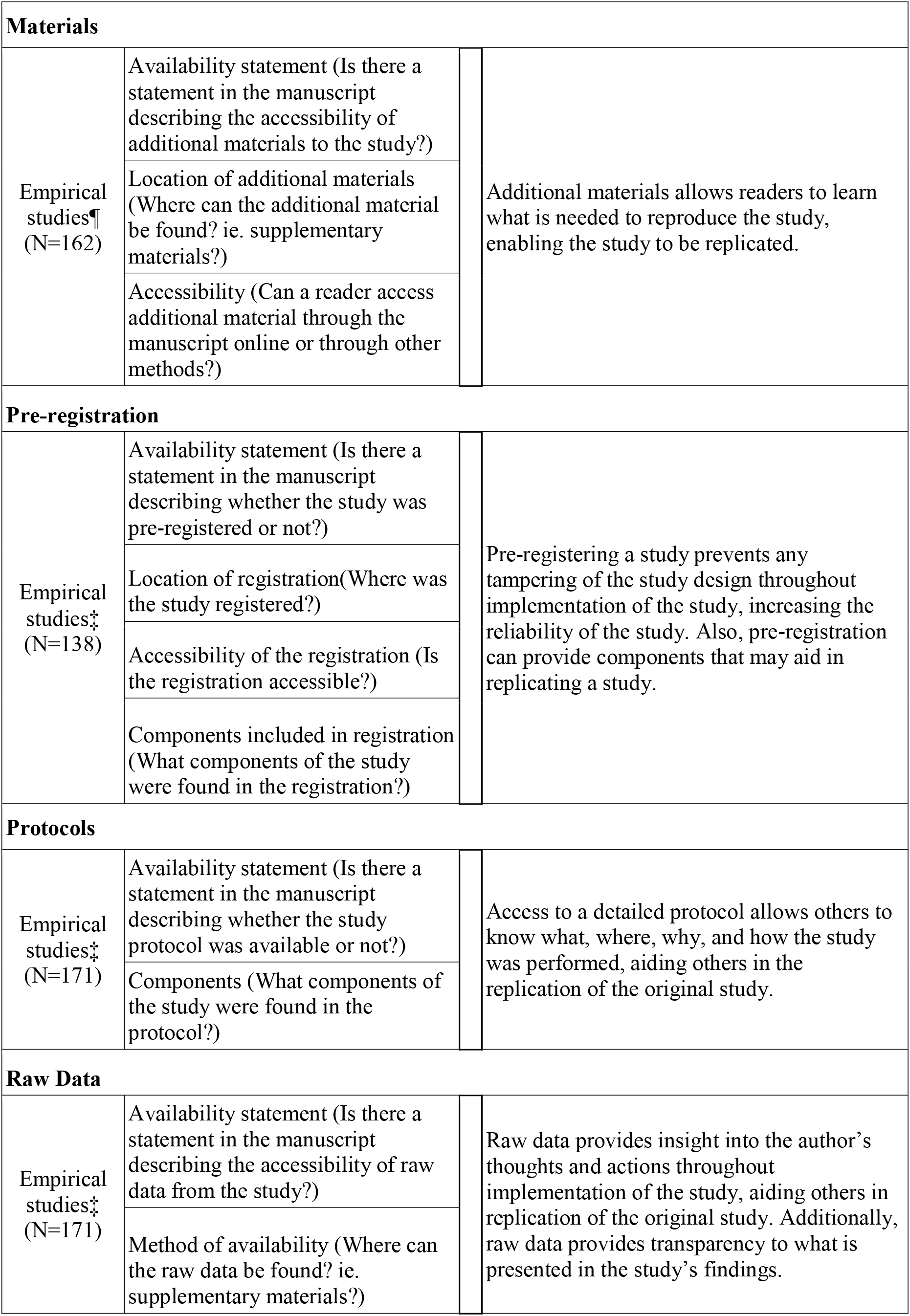
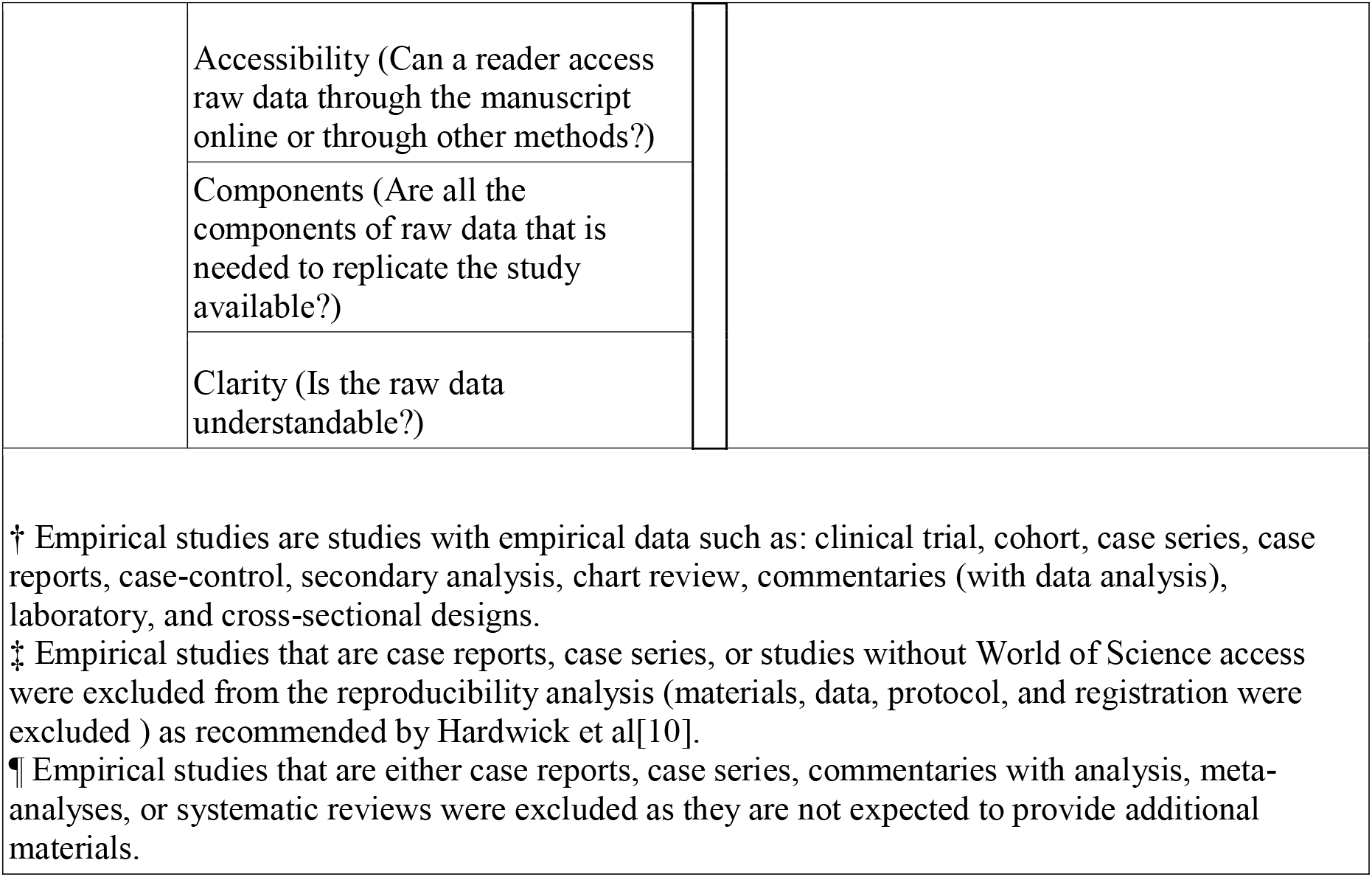
Types of Characteristics Associated with Reproducibility. Sample Sizes (N) depend on study type. Protocol about our measured characteristics is found online. (https://osf.io/x24n3/)

### Evaluation of Open Access Status

We evaluated all 300 publications to determine whether they were freely available online through open access. We searched Open Access Button (openaccessbutton.org) with publication titles and DOI numbers. This tool actively searches for the full-text online. If it could not find a publication, two of us (S.R. and B.J.) searched Google Scholar and PubMed to determine if the full text was available via open access on the journal website.

### Evaluation of Replication and Whether Publications Were Included in Research Synthesis

For empirical studies, excluding meta-analysis and commentary with analysis, we searched the Web of Science to determine whether the publication was cited in a replication study, meta-analysis, or systematic review. The Web of Science additionally lists information important for our study, such as the country of journal publication, 5-year impact factor (when available), and most recent impact factor.

### Statistical Analysis

We report descriptive statistics for each of our findings with 95% confidence intervals (95% CIs) using analysis functions within Microsoft Excel.

## Results

### Included Sample and Characteristics

Our inclusion criteria resulted in 42,422 articles from 46 urology journals found in the NLM catalog. Of the articles meeting the inclusion criteria, 300 articles were randomly chosen for analysis. Six articles were not analyzed because we did not have access to the text. The remaining 294 articles were assessed to determine the 5-year impact factor of their corresponding journals. Twenty of the 294 articles came from journals without 5-year impact factors. Thus, journals of the 274 studies reported a median of 2.466 as their 5-year impact factor with an interquartile range of 1.898 to 4.925. In addition, a full assessment of the original 300 articles revealed that 88 (29.33%) were accessible through Open Access Button or other means. Over half of our included studies (163/294, 55.44%) provided a statement revealing that their study was without a conflict of interest. However, 95 (32.31%) of our included studies did not provide any type of conflict of interest statement. Nearly two-thirds of our studies (185, 62.93%) did not state if or from where they received funding. Among the 109 studies that provided a statement, most did not receive funding (31, 28.44%). Of the 78 studies that did receive funding, most obtained it through public entities (23, 29.49%). Other characteristics of our included studies can be found in Table 2 and Supplementary Table 1.

**Table 2:** Characteristics of Included Publications

| Table 2: Characteristics of Included Publications |  |  |
| --- | --- | --- |
| Characteristics |  | Variables |
|  |  | N (%) |
| Funding<br>(N=294) | University | 4 (1.36%) |
|  | Hospital | 1 (0.34%) |
|  | Public | 23 (7.82%) |
|  | Private/Industry | 20 (6.80%) |
|  | Non-Profit | 2 (0.68%) |
|  | Mixed | 28 (9.52%) |
|  | No Statement Listed | 185 (62.93%) |
|  | No Funding Received | 31 (10.54%) |
| Type of Study<br>(N=294) | No Empirical Data | 115 (39.12%) |
|  | Meta-Analysis | 9 (3.06%) |
|  | Chart Review | 10 (0.34%) |
|  | Clinical Trial | 22 (7.48%) |
|  | Case Study | 6 (2.04%) |
|  | Case Series | 2 (0.68%) |
|  | Cohort | 94 (31.97%) |
|  | Case Control | 2 (0.68%) |
|  | Survey | 8 (2.72%) |
|  | Laboratory | 17 (5.78%) |
|  | Other | 9 (3.06%) |
| 5 Year Impact<br>Factor<br>(N=274) | Median | 2.466 |
|  | 1st Quartile | 1.898 |
|  | 3rd Quartile | 4.925 |
|  | Interquartile Range | 1.898 - 4.925 |

### Characteristics Associated with Reproducibility

The only studies that were assessed for reproducibility were those that had empirical data. Thus, 115 articles without empirical data were excluded from the initial 294 studies. We also excluded eight case studies and case series because such studies cannot be reproduced. We therefore assessed a total of 171 studies for reproducibility. Of these studies, 163 (95.32%) did not provide a pre-registration statement. Among the 8 studies that provided a pre-registration statement, 4 had accessible links to the pre-registration. Nearly all analyzed studies omitted a data availability statement (162/171, 94.74%). Of the 9 studies that provided a data statement, 2 claimed that their data was not available. None of the 7 studies that claimed their data were available provided enough raw data for the study to be reproduced. Similarly, 156 (96.30%) of 162 analyzed studies (excluding meta-analyses) did not provide a material availability statement. Six studies provided a material availability statement; five of these publications included a statement that materials were available, but only four provided working links to the materials. Only one of the 171 studies included a full protocol in the publication, and none of the 171 studies provided an analysis script availability statement. More characteristics associated with reproducibility are presented in Supplementary Table 1.

## Discussion

Our study revealed concerning findings regarding the reproducibility of research in urology literature. Only nine studies made statements regarding the availability of data, and only seven of those actually made their data available. Fewer than half of the studies in our sample were available through Open Access Button, and detailed protocols and pre-registration were rare. One trial in our sample was claimed to be a replication of a previous study, but even this publication failed to include any of the reproducibility markers that we assessed. These findings are similar to those of Hardwicke *et al[10]* for a survey of reproducibility in social sciences.

Our study revealed that only one study contained a link to protocols, while no studies provided analysis scripts and only six provided materials statements. These elements are the three most important ones in reproducing a study. Protocols provide details about how each step of the study was performed, to an extent much deeper than would be relevant to the average person reading the methods section.[13,14] Similarly, analysis scripts are crucial for re-creating the original analysis in a stepwise manner.[15] Materials include anything that was necessary for the study to be performed, including forms, questionnaires, devices, software programs, and more.[16] Some investigators have posited that freely providing these elements invites plagiarism of study design, a major concern with the pressure on researchers to publish while limiting time and funding.[17] Chan *et al[18]* have suggested placing protocols in a lockbox and making them available upon data release to protect intellectual property, while maintaining reproducible research. At the very least, authors should state in their articles that these crucial elements of reproducibility are available upon reasonable request.

Pre-registration is one of the best ways to increase transparency and reproducibility in research, yet only eight studies from our sample were pre-registered. Pre-registration of trials encourages transparency in research by outlining the intended outcomes, interventions, protocols, and methods of analysis before the study is underway.[19] When trials are not pre-registered, investigators have the freedom to manipulate data to obtain significance (P-hacking)[20], hypothesize after results are known (HARKing)[21], switch primary outcomes[22], or deviate from a priori protocols.[23] Several researchers, including Nosek *et al[24]* have called for widespread adoption of pre-registration, citing its value in increasing transparency, rigor, and reproducibility. Early results of pre-registration are positive, with pre-registered studies exhibiting a significant increase in null findings.[25] The OSF hosts pre-registration free of charge and also provides pre-registration templates and instructional guides.[26,27] High-impact journals could require pre-registration for any study to be considered for publication, which would encourage authors to take the necessary steps to increase the chance of having their research published in a respected journal.

Data availability is another area in which urology research falls short. Some journals, including *European Urology*, have begun to require that authors’ manuscripts include a description of how readers can access underlying data, while other journals mandate the inclusion of study protocols, analysis scripts, and any other element needed to replicate the original study.[28,29] Beginning in 2019, the International Committee of Medical Journal Editors (ICMJE) mandated data sharing statements by all prospective clinical trials submitted for publication to an ICMJE member journal.[30] Showing that such policies can be successful, *PLoS One*, another journal requiring data availability, reported that 20% of studies published in the journal hosted their data on a third-party website, 60% provided their data in a supplement, and the remaining 20% made their data available upon reasonable request.[31] These initiatives are steps in the right direction, and we propose a few more possibilities for improving reproducibility in urology research.

The Repeat framework was designed by McIntosh *et al[32]* to improve reproducibility in research. This easy-to-use checklist can be adapted for most studies. Additionally, the OSF developed the Transparency and Openness Promotion (TOP) Guidelines, which provide eight modular standards designed to increase transparency, disclosure, openness, and collaboration.[33] The EQUATOR network has set out to improve research reporting and manuscript writing through the use of reporting guidelines.[34,35] These guidelines, available for nearly every type of study, ensure that manuscripts are written in a transparent way, encouraging reproducibility and accurate reporting of findings.[12] Some journals have begun to require the use of reporting guidelines in the studies they publish.[36–38]

Our study has both strengths and limitations. Regarding strengths, we applied double data extraction procedures, which is considered a best practice methodology by the systematic review community and is recommended in the *Cochrane Handbook for Systematic Reviews of Interventions*.[44] To foster study reproducibility and transparency, we have made all relevant study materials publicly available on OSF. Concerning limitations, our study is cross-sectional in nature, including only PubMed-indexed journals that were published in English during a finite time period. Thus, our results should be interpreted in light of these considerations. Additionally, many replication studies are not published because they are never submitted for publication.[2] In recent years, some organizations, including Elsevier, have encouraged the submission and publication of replication studies, but they are not yet common in biomedical literature.[45] We did not attempt to contact authors for data availability, analysis scripts, protocols, or any of the other markers of reproducibility. While we may have found these things to be readily available, it is more likely that we would have run up against the familiar issues of low response rate and limited cooperation.[46,47]

## Conclusion

Current urology research does not consistently provide the components needed to reproduce original studies. Collaborative efforts from investigators and journal editors are needed to improve research quality, while minimizing waste and patient risk.

## Conflicts of Interest and Source of Funding

This study was funded through the 2019 Presidential Research Fellowship Mentor–Mentee Program at Oklahoma State University Center for Health Sciences.

---

**Supplemental 1:** Additional Characteristics of Reproducibility in Urology Studies

|  |  |  |  |
| --- | --- | --- | --- |
| **Characteristics** |  | **Variables** |  |
|  |  | **N (%)** | **95% CI** |
| **Conflict of Interest Statement (N=294 )** | Statement, one or more conflicts of interest | 36 (12.24%) | 8.54 - 15.95 |
|  | Statement. no conflict of interest | 163 (55.44%) | 49.82 - 61.07 |
|  | No conflict of interest statement | 95 (32.31%) | 27.02 - 37.61 |
| **Data Availability (N=171)** | Statement, some data are available | 7 (4.09%) | 1.85 - 6.34 |
|  | Statement, data are not available | 2 (1.17%) | 0 - 2.39 |
|  | No data availability statement | 162 (94.74%) | 92.21 - 97.26 |
| **Material Availability (N=162)** | Statement, some materials are available | 5 (3.09%) | 1.13 - 5.04 |
|  | Statement, materials are not available | 1 (0.62%) | 0 - 1.50 |
|  | No materials availability statement | 156 (96.30%) | 94.16 - 98.43 |
| **Protocol Availability (N=171)** | Full Protocol | 1 (0.58%) | 0 - 1.45 |
|  | No Protocol | 170 (99.42%) | 98.55- 100 |
| **Analysis Scripts (N= 171)** | Statement, some analysis scripts area vailable | 0 (0%) | - |
|  | Statement, analysis scripts are not available | 0 (0%) | - |
|  | No analysis script availability statement | 171 (100%) | - |
| **Replication Studies (N= 171)** | Novel study | 170 (99.42%) | 98.55 - 100 |
|  | Replication | 1 (0.58%) | 0 - 1.45 |
| **Open Access (N= 300)** | Yes - found via Open Access Button | 87 (29.00%) | 23.87 - 34.13 |
|  | Yes - found article via other means | 1 (0.33%) | 0 - 0.99 |
|  | Could not access through paywall | 212 (70.67%) | 65.51 - 75.82 |
| **Cited in Systematic Review/ Meta-Analysis (a) (N=169)** | No Citations | 140 (82.84%) | 78.57 - 87.11 |
|  | A Single Citation | 20 (11.83%) | 8.18 - 15.49 |
|  | One to Five Citations | 9 (5.33%) | 2.78 - 7.87 |
| Abbreviations: CI, Confidence Interval. a - No studies were explicitly excluded from the systematic reviews or meta-analyses that cited the original article. |  |  |  |
| **Pre-Registration (N= 171)** | Statement, says was pre--registered | 8 (4.68%) | 2.29 - 7.07 |
|  | Statement, says was not pre-registered | 0 | - |
|  | No, there is no pre-registration statement | 163 (95.32%) | 92.93 - 97.71 |
| **Test Subjects (N= 294)** | Animals | 11 (3.74%) | - |
|  | Humans | 204 (69.39%) | - |
|  | Neither | 79 (26.87%) | - |
| **Country of Journal Publication (N=294)** | US | 182 (61.90%) | - |
|  | UK | 34 (11.56%) | - |
|  | Germany | 2 (0.68%) | - |
|  | India | 11 (3.74%) | - |
|  | Italy | 3 (1.02%) | - |
|  | Unclear | 9 (3.06%) | - |
|  | Other (a) | 53 (18.03) | - |
| **Country of Corresponding Author (N=294)** | US | 116 (39.46%) | - |
|  | China | 20 (6.80%) | - |
|  | UK | 14 (4.76%) | - |
|  | Germany | 10 (3.40%) | - |
|  | Japan | 13 (4.42%) | - |
|  | France | 6 (2.04%) | - |
|  | Canada | 14 (4.76%) | - |
|  | Italy | 24 (8.16%) | - |
|  | India | 2 (0.68%) | - |
|  | Spain | 5 (1.70%) | - |
|  | Unclear | 11 (3.74%) | - |
|  | Other | 59 (17.69%) | - |

**Supplemental 2:**
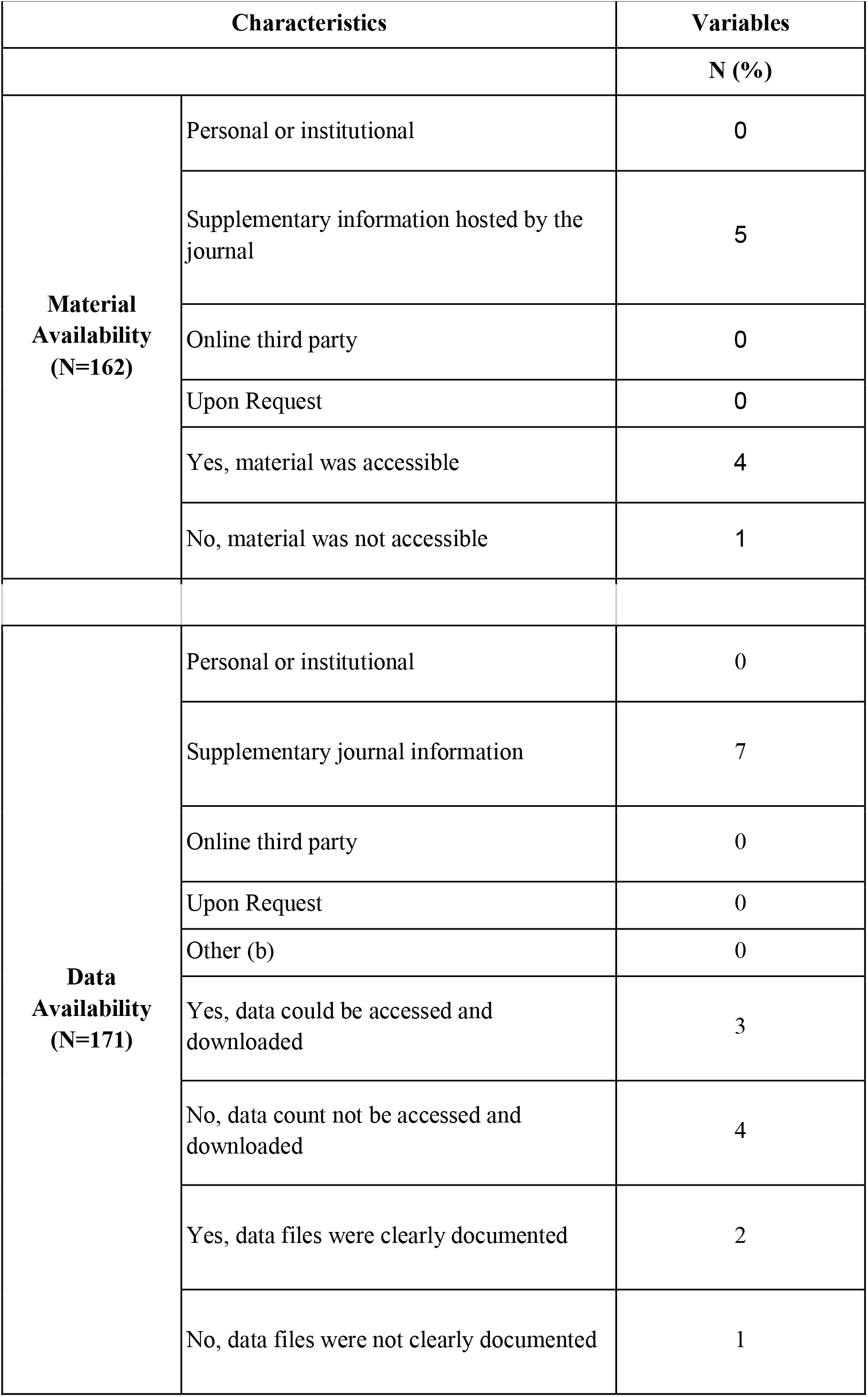
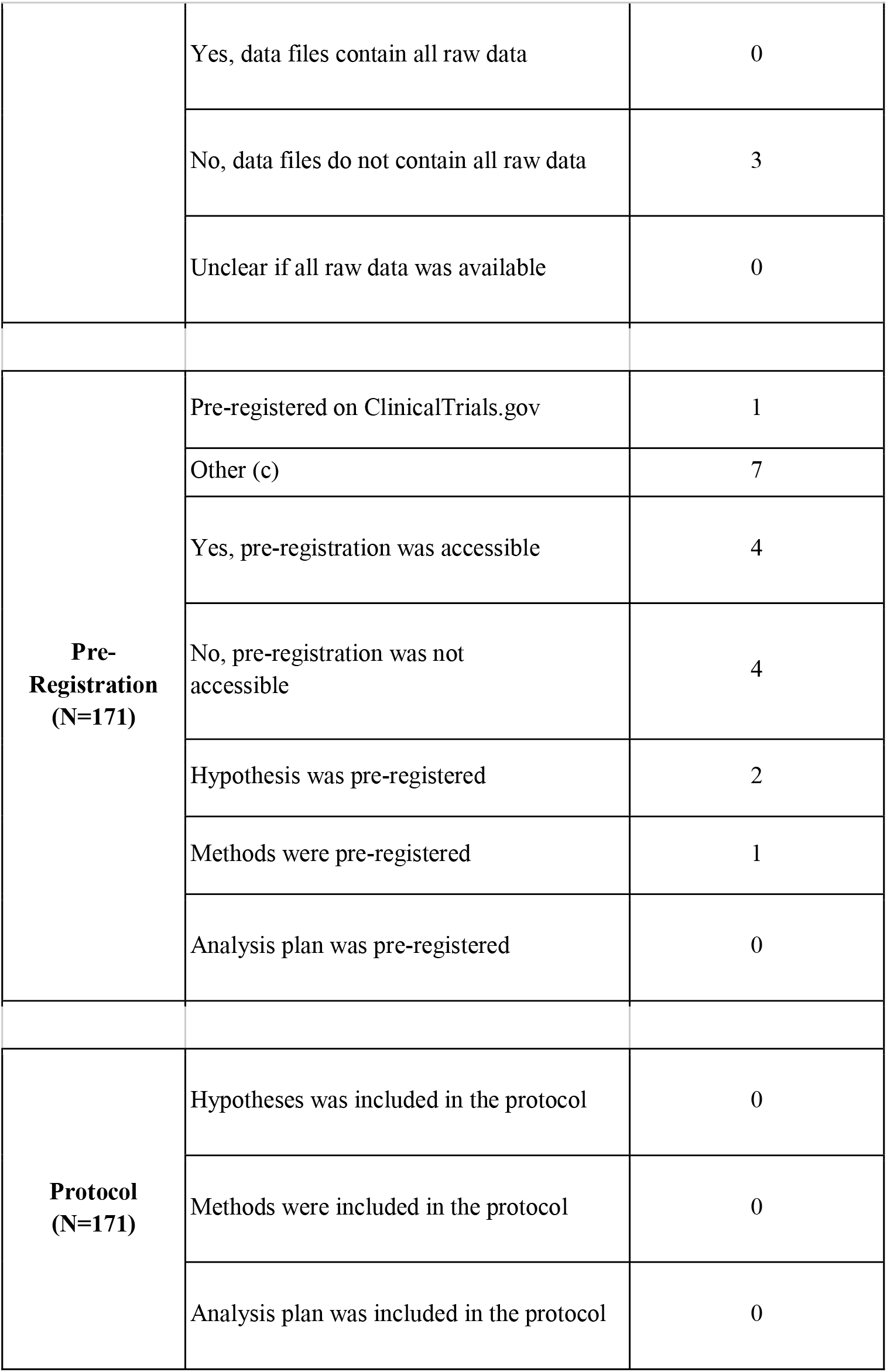
Additional Characteristics of Reproducibility in Urology Studies

|  |  |  |
| --- | --- | --- |
| **Characteristics** |  | **Variables** |
|  |  | **N (%)** |
| **Material Availability (N=162)** | Personal or institutional | 0 |
|  | Supplementary information hosted by the journal | 5 |
|  | Online third party | 0 |
|  | Upon Request | 0 |
|  | Yes, material was accessible | 4 |
|  | No, material was not accessible | 1 |
| **Data Availability (N=171)** | Personal or institutional | 0 |
|  | Supplementary journal information | 7 |
|  | Online third party | 0 |
|  | Upon Request | 0 |
|  | Other (b) | 0 |
|  | Yes, data could be accessed and downloaded | 3 |
|  | No, data could not be accessed and downloaded | 4 |
|  | Yes, data files were clearly documented | 2 |
|  | No, data files were not clearly documented | 1 |
|  | Yes, data files contain all raw data | 0 |
|  | No, data files do not contain all raw data | 3 |
|  | Unclear if all raw data was available | 0 |
| **Pre-Registration**<br>**(N=171)** | Pre-registered on ClinicalTrials.gov | 1 |
|  | Other (c) | 7 |
|  | Yes, pre-registration was accessible | 4 |
|  | No, pre-registration was not accessible | 4 |
|  | Hypothesis was pre-registered | 2 |
|  | Methods were pre-registered | 1 |
|  | Analysis plan was pre-registered | 0 |
| **Protocol**<br>**(N=171)** | Hypotheses was included in the protocol | 0 |
|  | Methods were included in the protocol | 0 |
|  | Analysis plan was included in the protocol | 0 |

